# A novel meiotic drive in Neurospora crosses heterozygous for hybrid translocation strains disfavors homokaryotic progeny derived from alternate, but not adjacent-1 segregation

**DOI:** 10.1101/039487

**Authors:** Dev Ashish Giri, S. Rekha, Durgadas P. Kasbekar

## Abstract

Four insertional or quasiterminal translocations (*T*) were recently introgressed from Neurospora crassa into *N. tetrasperma*. Crosses of two of the resulting *T*^*Nt*^ strains with *N. tetrasperma N strains* (*N* = normal sequence) produced more *Dp* than *T* and *N* homokaryotic progeny, although [*T* + *N*] and [*Dp* + *Df*] heterokaryotic progeny were made in roughly equal numbers. The *T*, *N*, and [*T* + *N*] progeny are derived from alternate segregation (ALT), whereas adjacent-1 segregation (ADJ) generates the *Dp*, *Df*, and [*Dp* + *Df*] types. Differential recovery of homokaryotic products from ALT and ADJ represents a novel and unprecedented type of meiotic drive. This drive contributed to our inability to introgress a larger insertions translocation, *T*(*VR*>*VIL*)*UK3-41*, into *N. tetrasperma.* We suggest that one or more Bateson-Dobzhansky-Muller type incompatibility between *N. crassa* and *N. tetrasperma* genes in the *T*^*Nt*^ x *N* crosses might cause an insufficiency for a product required for ascospore maturation. Since the *Df* type is inviable, only four ascospores (*Dp* or [*Dp* + *Df*] types) share this limited resource in [*Dp* + *Df*] asci, whereas four to eight ascospores compete for it in [*T* + *N*] asci. This increases the chance that in asci with >4 ascospores none properly matures, and results in *Dp* progeny out-numbering *T* and *N* types.

## Introduction

Recently, we introgressed four insertional translocations (*IT*; EB4, IBj5, UK14-1, and B362i) from *Neurospora crassa* into *N. tetrasperma*, to make heterokaryons with complementary duplications and deficiencies in their constituent nuclei ([*Dp* + *Df*]; Giri *et al*., 2015). Construction of the [*Dp* + *Df*] heterokaryon relied on (1) the difference in *N. crassa* and *N. tetrasperma* ascogenesis, and (2) adjacent-1 segregation in *IT*-heterozygous crosses. The introgressed strains, designated as *T(EB4)*^*Nt*^; *T(IBj5)*^*Nt*^; *T(UK14-1)*^*Nt*^ and *T(B362i)*^*Nt*^, nominally have a *N. tetrasperma*-derived genome, except at the *N. crassa*-derived breakpoint junctions. Now, we report that crosses of some of these *T*^*Nt*^ strains with normal sequence strains (i.e., *T*^*Nt*^ x *N*) exhibit a novel type of meiotic drive that targets the homokaryotic progeny made following alternate, but not adjacent-1, segregation.

In Neurospora sexual crosses, fusion of haploid *mat A* and *mat a* nuclei produces a diploid zygote nucleus that undergoes meiosis and a post-meiotic mitosis. In *N. crassa*, the eight haploid progeny nuclei (4 *mat A* + 4 *mat a*) are partitioned into the eight initially uninucleate ascospores formed per ascus, whereas in *N. tetrasperma* the asci make four initially binucleate ascospores, each receiving a pair of non-sister nuclei (1 *mat A* + 1 *mat a*), although occasionally, a pair of smaller homokaryotic ascospores replaces one or more dikaryotic ascospore (Raju 1992; Raju and Perkins 1994). The dominant *Eight-spore* (*E*) mutant increases such replacement, and can generate asci with upto eight ascospores (Dodge, 1939; Calhoun and Howe, 1968). The homokaryotic *N. crassa* mycelia, of *mat A* or *mat a* mating type, can mate only with mycelia from another ascospore of the opposite mating type, whereas the dikaryotic [*mat A* + *mat a*] *N. tetrasperma* mycelia can undergo a self-cross. Dikaryotic *N. tetrasperma* mycelia also produce some homokaryotic conidia (vegetative spores), and the mycelia produced from homokaryotic conidia and ascospores can out-cross with like mycelia of the opposite mating type (Raju and Perkins 1994; Bistis, 1996).

Perkins (1997) described three types of translocations (*T*) in *N. crassa*, viz, insertional (*IT*), quasiterminal (*QT*) and reciprocal (*RT*). An *IT* transfers a segment from a donor chromosome into a recipient chromosome, without any reciprocal exchange, and creates three breakpoint junctions; A on the donor chromosome, and B and C (proximal and distal) on the recipient chromosome (Singh *et al*., 2010). *QT*s transfer a donor chromosome’s distal segment to the tip of a recipient chromosome distal to any essential gene, and presumably the recipient chromosome’s tip caps the donor chromosome’s break; and *RT*s reciprocally interchange the terminal segments of two chromosomes. *QT*s and *RT*s create two breakpoint junctions. Crosses of *IT*s and *QT*s with *N* (*N* = normal sequence) can produce *T*, *Dp*, and *N* progeny (Perkins, 1997). *T* progeny contain the A, B, and C breakpoints, *Dp* contain B and C but not A, and *N* contain none. Breakpoint junctions of several *T*s have previously been determined (Singh 2010; Singh *et al*. 2010).

In *T* x *N* crosses homologous centromeres segregate either by alternate (ALT) or adjacent-1 (ADJ) segregation. In *N. crassa*, ALT produces eight parental-type viable black (B) ascospores (4*T* + 4*N*), whereas ADJ produces eight non-parental-type ascospores, which in *IT* x *N* and *QT* x *N* include four viable black duplication ascospores and four inviable white (W) ones with the complementary deficiency (4 *Dp* + 4 *Df*) (Perkins, 1997). Since ALT and ADJ are equally likely, the crosses produce equal numbers of 8B:0W and 4B:4W asci. Additionally, recombination in the interstitial regions between the centromeres and breakpoints can generate 6B:2W asci, but 2B:6W and 0B:8W types are not expected (Perkins 1997). In contrast, isosequential crosses (i.e., *N* x *N* or *T* x *T*) produce mostly 8B:0W asci (Perkins 1997).In *RT* x *N*, all eight ascospores from ADJ have complementary *Dp*/*Df* (i.e., *Dp2*/*Df1* and *Dp1*/*Df2*), and the asci are 0B:8W, thus 8B:0W = 0B:8W is characteristic of *RT* x *N*.

ALT and ADJ in *IT* (and *QT*) x *N* crosses in *N. tetrasperma*, generate 4B:0W asci with, respectively, [*T* + *N*] and [*Dp* + *Df*] ascospores (Giri *et al*., 2015). The two genotypes can inter-convert via self-crosses. The RIP mutational process, targeting duplicated DNA (Selker 1990), can occur in self-crosses of [*Dp* + *Df*] but not [*T* + *N*]. Crosses of *T*^*Nt*^ strains with *N. tetrasperma* strains 85 *A* or 85 *a* produced mostly four-spored asci, but the rare eight-spored asci included 8B:0W, 4B:4W, and 6B:2W types (Giri *et al*., 2015). More eight-spored asci were produced in crosses of *T*^*Nt*^ with *E*. The crosses of *T(EB4)*^*Nt*^ and *T(UK14-1)*^*Nt*^ with *E* produced comparable numbers of 8B:0W and 4B:4W asci, and also 6B:2W. Surprisingly, the crosses of *T(IBj5)*^*Nt*^ and *T(B362i)*^*Nt*^ with *E* yielded no 8B:0W and 6B:2W asci, suggesting that maturation of eight-spored post-ALT asci, but not post-ADJ asci, is blocked (Giri *et al*., 2015). Since no ascus contained more than four black ascospores we call this the max-4^-^ (m^-^) phenotype (Figure 1). A model was proposed based on the facts that (1) the *EA, Ea*, and *C4,T4 a* strains are all derived from the *N. tetrasperma* strain 343.6 *A E* (Metzenberg and Ahlgren 1969), and (2) the *C4,T4 a* strain was used for *T*^*Nt*^ construction. In this model (Giri *et al*., 2015) the *E*, *T*(*IBj5*)^*Nt*^, and *T*(*B362i*)^*Nt*^ strains share a 343.6 *A E*- derived recessive mutation that blocks maturation of post-ALT eight-spored asci, and the mutation is absent from 85 *A*, 85 *a*, *T(EB4)*^*Nt*^ and *T*(*UK14-1*)^*Nt*^. Homozygosity in *T(IBj5)^Nt^ a* × *E A* and *T(B362i)^Nt^ A* × *E a* produces m^-^ asci, whereas heterozygosity in *T(EB4)^Nt^ a* × *E A* and *T(UK14-1)^Nt^ A* × *E a* produces normal asci. We now show that this model is not tenable. Instead the m^-^ phenotype appears to reflect a meiotic drive that specifically disfavors the homokaryotic progeny made following ALT.

**Figure 1.**
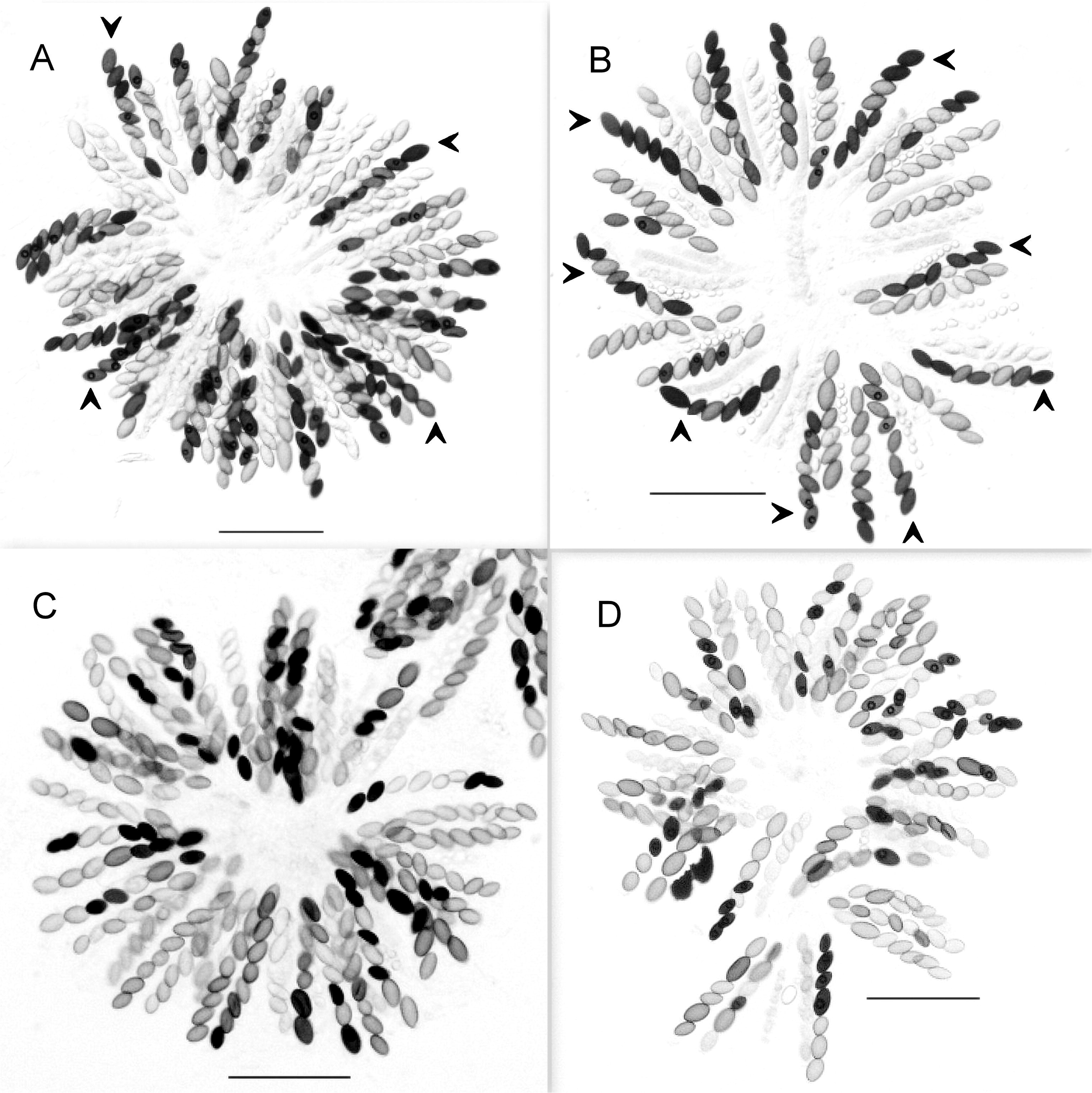
The max-4^-^ (m^-^) phenotype. Asci from the cross *T(EB4)*^*Nt*^ *a* x *E a* are shown in the upper panel, and from *T(B362i)*^*Nt*^ *A* x *E a* in the lower panel. Arrowheads in the upper panel identify some asci with more than four black ascospores. No ascus in the lower panel has more than four black ascospores. This defines the m^-^ phenotype. The black ascospores are presumed to be viable and the white ones inviable. The scale-bar in all images is 100 ìm.

## Materials and methods

### 1. Neurospora strains and general genetic manipulations

Neurospora genetic analysis was done essentially as described by Davis and De Serres (1970). Metzenberg’s (2003) alternative recipe was used for making Medium N. The *N. crassa T*(*VR*>*VIL*)*UK3-41*, *inl A* strain (FGSC 6869) was obtained from the Fungal Genetics Stock Center (FGSC, Department of Plant Pathology, Kansas State University), as were the standard *N. tetrasperma* strains 85 *A* (FGSC 1270) and 85 *a* (FGSC 1271); the *E* mutants *lwn; al(102)*, *E A* (FGSC 2783) and *lwn; al(102)*, *E a* (FGSC 2784) (hereafter *E A* and *E a*), and the *N. crassa* / *N. tetrasperma* hybrid strain: *C4,T4 a* (FGSC 1778). The *C4,T4 a* strain has four *N. crassa* great-grandparents and four *N. tetrasperma* great-grandparents (Metzenberg and Ahlgren 1969). The *N. crassa* great-grandparents were of the OR background, whereas the *N. tetrasperma* great-grandparents were of the 343.6 *A E* background (Metzenberg and Ahlgren 1969).

The *T(EB4)*^*Nt*^ *a*; *T(IBj5)*^*Nt*^ *a*; *T(UK14-1)*^*Nt*^ *A* and *T(B362i)*^*Nt*^ *a* strains were constructed by Giri *et al*, (2015) and have been deposited in the FGSC with the strain numbers FGSC 25016, FGSC 25017, FGSC 25018, and FGSC 25019. Briefly, the *N. crassa IT* strains *T(EB4)A*; *T(IBj5)A*; and *T(B362i)A,* and *QT* strain *T(UK14-1)A* (Perkins, 1997; Singh, 2010), were crossed with *C4T4 a* (Metzenberg and Ahlgren 1969), and the *T* progeny from these crosses were either again crossed with *C4T4* a, or with the opposite mating type derivative of *N. tetrasperma* 85 (i.e., 85 *A* or 85 *a*), and this step was reiterated until a *T* x 85 cross yielded self-fertile heterokaryotic progeny with all the three breakpoint junctions characteristic of the *T*. Such heterokaryons generally were [*T* + *N*] or [*Dp* + *Dp*] in genotype. Heterokaryotic [*T* + *N*] mycelia yield homokaryotic (self-sterile) conidial derivatives of both mating types, whereas [*Dp* + *Df*] mycelia yield homokaryons of only the mating type of the *Dp* nucleus. The *T*^*Nt*^ strains were isolated as self-sterile conidial derivatives from the [*T* + *N*] mycelia.

### 2. Markers polymorphic between the *85*/ *EA*/ *Ea* and FGSC 2508 *A*/FGSC 2509*a* strains

The genome sequence of the *N. tetrasperma* strain FGSC 2508 *A* and FGSC 2509 *a* is publicly available (fungidb.org). The *N. tetrasperma* strains *85*, *EA*, and *Ea* share considerable DNA sequence homology (data not shown), and their genome is more diverged from that of FGSC 2508*A* and FGSC 2509*a*. The oligonucleotide primers used for PCR and restriction enzymes used to digest the resulting amplicons to obtain polymorphisms between the *85*/*EA*/*Ea* and FGSC 2508 *A*/FGSC 2509 *a* alleles on each of the seven *N. tetrasperma* chromosomes are listed in Supplementary Table 1, and Supplementary Figure 1 shows the marker positions on the chromosomes.

### 3. PCR-based identification of *T*, *N*, and *Dp* progeny from *T*^*Nt*^ x *N* crosses

*IT*s are defined by three breakpoint junctions; A on the donor chromosome, and B and C (proximal and distal) on the recipient chromosome, whereas *QT*s have two breakpoint junctions, A and B, on the two participating chromosomes. *T* progeny contain the A, B, and C breakpoints, *Dp* contain B and C but not A, and *N* contain none. As positive controls we used additional primers to amplify by PCR genome segments from the *N*-derived homologues of the donor and recipient chromosomes (designated N^D^ and N^R^) but not from the translocation chromosomes. The primers are listed in Supplementary Table 2.

The *N. crassa* insertional translocation *T(VR>VIL)UK3-41* translocates 1879356 bp, bearing 490 genes, from chromosome 5R to 6L (Perkins, 1997). Its translocated segment is larger than all the four previously introgressed translocations combined (Singh 2010; Singh *et al*., 2010). Its A breakpoint junction was previously determined (Genbank accession number HM573450), but the B and C breakpoint junction sequences were determined in this work. Genomic DNA of the *T(UK3-41)* strain was digested with *Hae* III, self-ligated, and the product used as template in inverse PCR reactions with the primers listed in Supplementary Table 3.

### 4. Introgression of *T*(*VI* > *V*)*UK3-41*

Crosses between standard *N. crassa* strains of the Oak Ridge (OR) background and *N. tetrasperma* strains of the 85 background are almost completely sterile (Perkins 1991; Giri *et al*., 2015). However, both strain OR *A* and strain 85 *A* can cross with the *N. crassa* / *N. tetrasperma* hybrid strain *C4,T4 a* and produce viable progeny allowing the *C4,T4 a* strain to be used as a bridging strain for the initial introgression crosses.

The *T(UK3-41)* strain was crossed with *C4T4 a* and *T* progeny from these crosses (designated *T*^*1xC4T4*^) were distinguished from their *Dp* and *N* siblings by PCR with A and C breakpoint junction-specific primers. Nominally, 50% of the genome of *T*^*1xC4T4*^ progeny is derived from the *C4*,*T4 a* parent. The *T*^*1xC4T4*^A strains were crossed with *C4,T4 a*, to obtain the *T*^*2xC4T4*^ progeny in a like manner. Crosses of *T*^*2xC4T4*^ with the opposite mating type derivative of strain *85* were productive, and their *T* progeny were designated as *T*^*1x85*^. Likewise, *T*^*1x85*^ x *85* yielded *T*^*2x85*^, etc. Ordinarily, after two to three iterations of the crosses with *85*, we could recover progeny ascospores that produced mycelium of dual mating specificity characteristic of *N. tetrasperma*. That is, the resulting mycelium could cross with both 85*A* and *a*, and it could also undergo a self-cross. However, as documented in the Results section, by the f_4_ generation the deviation from the expected Mendelian ratio in the progeny caused us to run out of *T* progeny to proceed with additional introgression crosses.

## Results

### 1. The m ^-^ phenotype is not caused by a recessive mutation, and *E* maps to chr. 6

To test whether the *E*, *T(IBj5)*^*Nt*^, and *T*(*B362i*)^*Nt*^ strains, but not 85 *A* and 85 *a*, contain a recessive mutation whose homozygosity in *T* x *N* crosses blocks the maturation of post-ALT eight-spored asci, we screened 103 and 58 f1 segregants from *E A* x 85 *a* and *E a* x 85 *A*, and found, respectively, 57 and 21 homokaryons (i.e., self-sterile), crossed them with *T(IBj5)*^*Nt*^ *a* or *T(B362i)*^*Nt*^ *A*, and examined the crosses for the max-4^-^ (m^-^) phenotype (i.e., no asci with >4 black ascospores, Figure 1). We expected that the progeny inheriting the *E*-derived recessive allele would show the m^-^ phenotype, and those inheriting the 85-derived wild-type allele would produce normal asci. Unexpectedly, all 78 f1 progeny showed the m^-^ phenotype (data not shown). This is possible if the chromosome bearing the *E*-derived mutant allele also exerts meiotic drive relative to its 85-derived homologue. However, we were unable to easily find polymorphic markers between the *E* and 85 strains to test if such indeed was the case.

Instead, we found several polymorphic markers between the *85*/*EA*/*Ea* strains on the one hand and the *N. tetrasperma* FGSC 2508*A*/FGSC 2509*a* strains on the other (see Materials and methods). We verified that these markers showed independent segregation in the homokaryotic progeny from the crosses *Ea* x 2508*A* and *EA* x 2509*a* (Table 1). Further, crosses of *T(IBj5)^Nt^a* and *T(B362i)*^*Nt*^*A* with 2508*A* and 2509*a* produced mostly four-spored asci, but the rare eight-spored asci included some 8B:0W, 4B:4W, and 6B:2W types (Table 2). This showed that the presumptive recessive mutation for the m^-^ phenotype is absent from the 2508*A* and 2509*a* strains. However, all homokaryotic f_1_ progeny examined from *Ea* x 2508*A* (N = 14) and *EA* x 2509*a* (N = 20) gave the m^-^ phenotype in crosses with *T*(*IBj5*)^*Nt*^ *a* or *T(B362i)*^*Nt*^ *A* (data not shown). Given that all chromosomes segregate independently in the f_1_ progeny, these results are incompatible with the idea that the m^-^ phenotype segregates with a specific chromosome from *EA* or *Ea*. Thus, the hypothesis that the m^-^ phenotype is caused by homozygosity for a recessive mutation is not tenable.

**Table 1:**
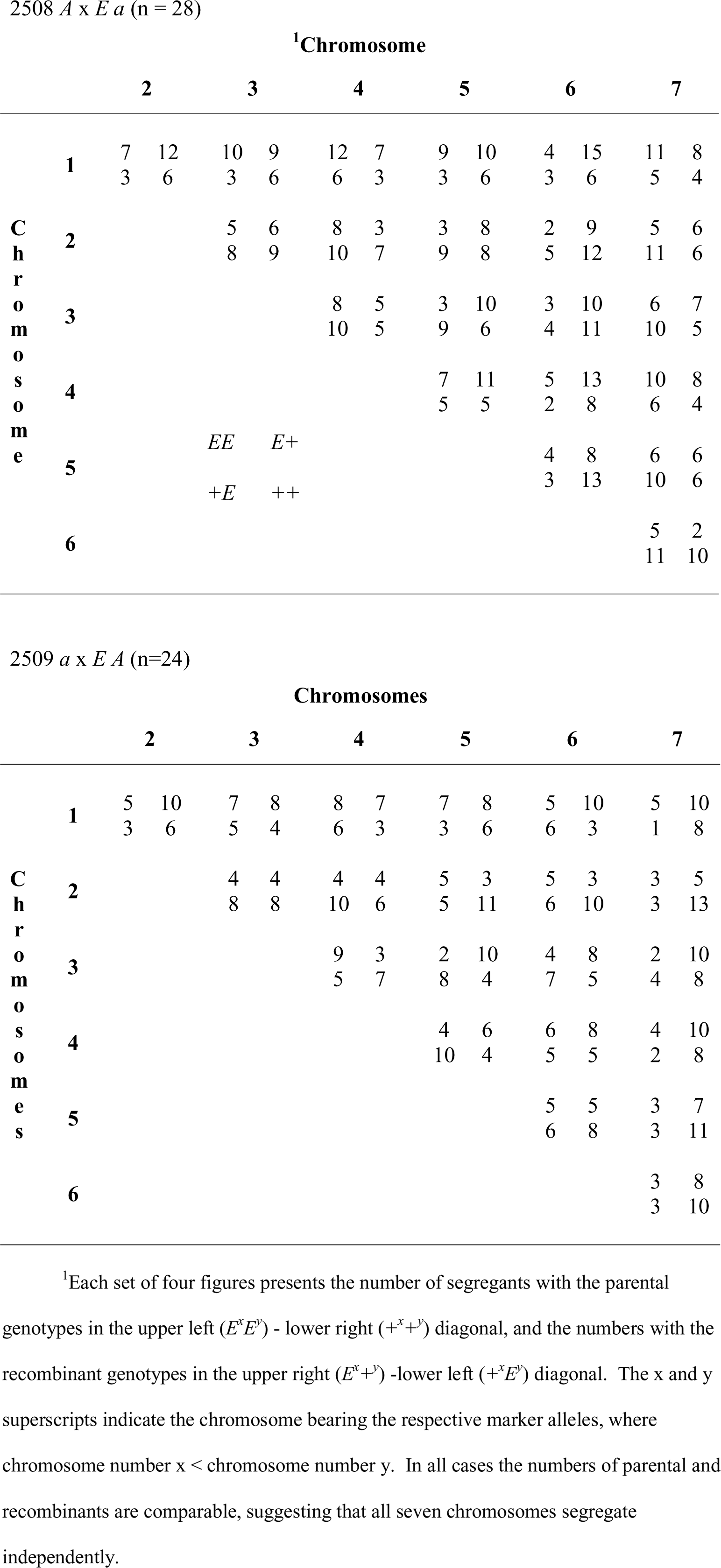
Segregation of chromosomes in the self-sterile progeny from the cross of *Ea* (or *EA*, *E*) with 2508*A* (or 2509*a*, +).

**Table 2:**
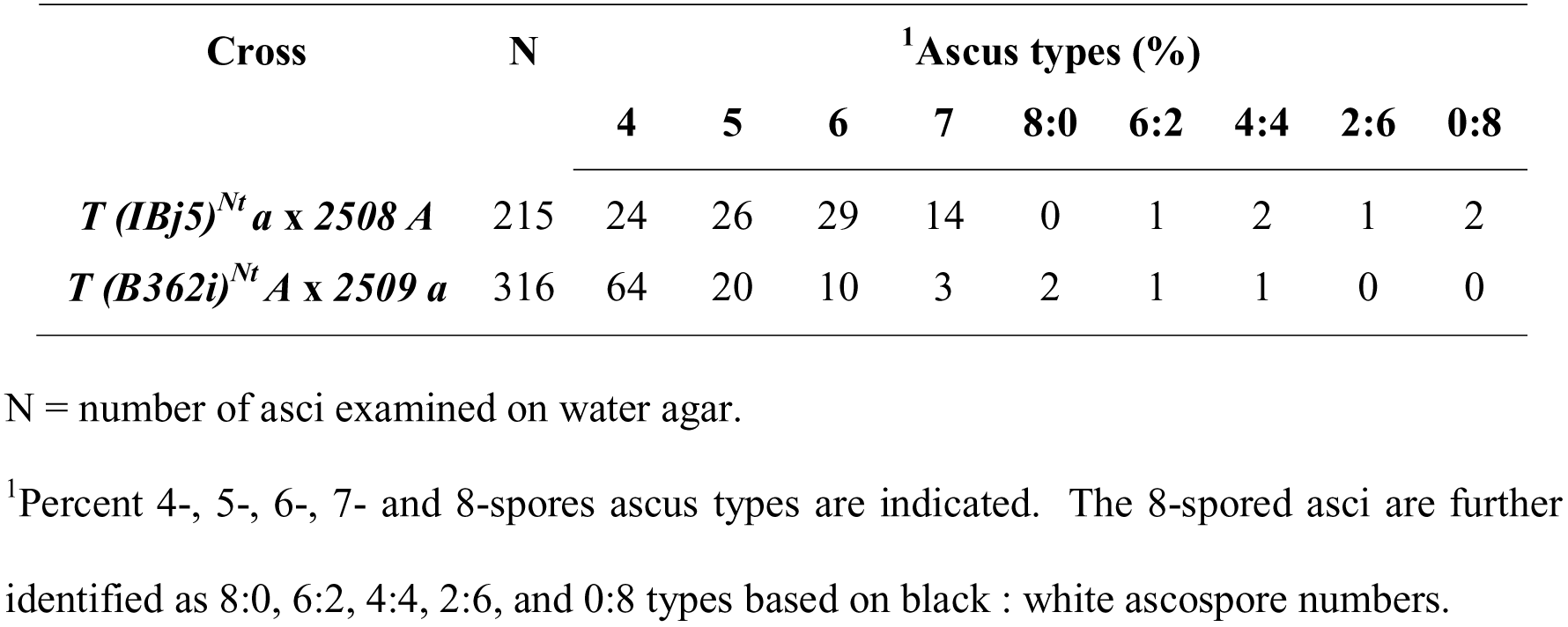
*T(IBj5)*^*Nt*^*a* and *T(B362i)*^*Nt*^*A* crossed with the normal sequence FGSC 2508 *A* or FGSC 2509 *a* strains yield eight-spored asci that include the 8B:0W and 6B:2W types.

| Cross | N | <sup>1</sup>Ascus types (%) |  |  |  |  |  |  |  |  |
| --- | --- | --- | --- | --- | --- | --- | --- | --- | --- | --- |
|  |  | 4 | 5 | 6 | 7 | 8:0 | 6:2 | 4:4 | 2:6 | 0:8 |
| $T (IBj5)^{Nt}a$ x 2508 A | 215 | 24 | 26 | 29 | 14 | 0 | 1 | 2 | 1 | 2 |
| $T (B362i)^{Nt}A$ x 2509 a | 316 | 64 | 20 | 10 | 3 | 2 | 1 | 1 | 0 | 0 |

N = number of asci examined on water agar.
<sup>1</sup>Percent 4-, 5-, 6-, 7- and 8-spores ascus types are indicated. The 8-spored asci are further identified as 8:0, 6:2, 4:4, 2:6, and 0:8 types based on black : white ascospore numbers.

The f_1_ homokaryotic segregants from *Ea* x 2508*A* and *EA* x 2509*a* were crossed with strain 85 derivatives of the opposite mating type and the progeny asci were examined for an eight- or four-spore phenotype. The results summarized in Table 3 show that the *E* mutation segregates independently of markers on all chromosomes, except chromosome 6. This confirms the mapping results of Howe and Haysman (1966). To localize *E* further we tested more f1 segregants and scored additional polymorphic loci on chromosome 6. The results of these studies will be presented elsewhere.

**Table 3:** The *Eight spore* mutation (*E*) segregates with chromosome 6.

|  |  | <sup>1</sup>Chromosomes |  |  |  |  |  |  |  |  |  |  |  |  |  |
| --- | --- | --- | --- | --- | --- | --- | --- | --- | --- | --- | --- | --- | --- | --- | --- |
|  |  | **1** |  | **2** |  | **3** |  | **4** |  | **5** |  | **6** |  | **7** |  |
|  |  | *E* | + | *E* | + | *E* | + | *E* | + | *E* | + | *E* | + | *E* | + |
| 2508 *A* x *E a*<br>(n = 25) | <sup>2</sup>**E** | 7 | 11 | 2 | 6 | 4 | 8 | 7 | 9 | 6 | 6 | 7 | 0 | 7 | 9 |
|  | <sup>2</sup>+ | 2 | 5 | 7 | 10 | 5 | 8 | 2 | 7 | 3 | 10 | 2 | 16 | 2 | 7 |
| 2509 *a* x *E A*<br>(n = 24) | **E** | 6 | 9 | 5 | 3 | 4 | 8 | 5 | 9 | 5 | 5 | 10 | 1 | 3 | 3 |
|  | + | 5 | 4 | 6 | 10 | 7 | 5 | 6 | 4 | 6 | 8 | 1 | 12 | 8 | 10 |

<sup>1</sup>(*E* +) = marker alleles from *E* and FGSC 2508 or FGSC 2509 strains. <sup>2</sup>E and <sup>2</sup>+ = Eight spored (**E**) or wild-type (+) ascus phenotypes. Each set of four figures presents the number of segregants with the parental **E** phenotype and *E* genotype or + phenotype and + genotype in the upper left (**EE**) - lower right (++) diagonal, and the numbers with the recombinant **E** phenotype and + genotype or + phenotype and *E* genotype in the upper right (**E+**) -lower left (+*E*) diagonal. The number of parental and recombinants segregants are comparable for markers on all chromosomes, except chromosome 6, for which parental > recombinants. This shows that the *E* mutation segregates with chromosome 6.

### 2. Deviation from the Mendelian ratio in progeny from some *T*^*Nt*^ x *N* crosses

We identified the self-sterile progeny (i.e., mating type homokaryons) from crosses of the *T*^*Nt*^ strains with *E* and 85 strains of the opposite mating type, and determined their *T*, *N*, or *Dp* genotype by PCR. As expected, and as can be seen in the results summarized in Table 4, more self-sterile progeny were produced from crosses with the *E* strains. In addition, we could discern three different phenotypes with regard to numbers of *T*, *N*, and *Dp* progeny: crosses of *T(EB4)*^*Nt*^*a* with 85 *A* and *E A* produced comparable numbers of *T*, *N*, and *Dp* progeny (*T* = *N* = *Dp*, phenotype 1), whereas *T(IBj5)*^*Nt*^*a* x *E A*, *T(IBj5)*^*Nt*^*a* x 85 *A*, and *T(B362i)*^*Nt*^*a* x 85 *a* gave *T* and *N* ≪ *Dp* (phenotype 2), and *T(B362i)*^*Nt*^*A* x *E a* gave *T* ≪ *N* and *Dp* (phenotype 3). For *T(UK14-1)*^*Nt*^*a* x *N A* only the *T* progeny were identifiable by their A breakpoint junction, whereas *Dp* and *N* could not be distinguished because the B junction of *T(UK14-1)*^*Nt*^, a *QT*, is not yet defined (Table 4). *Dp* have only the B breakpoint and not A, whereas *N* have neither.

**Table 4:**
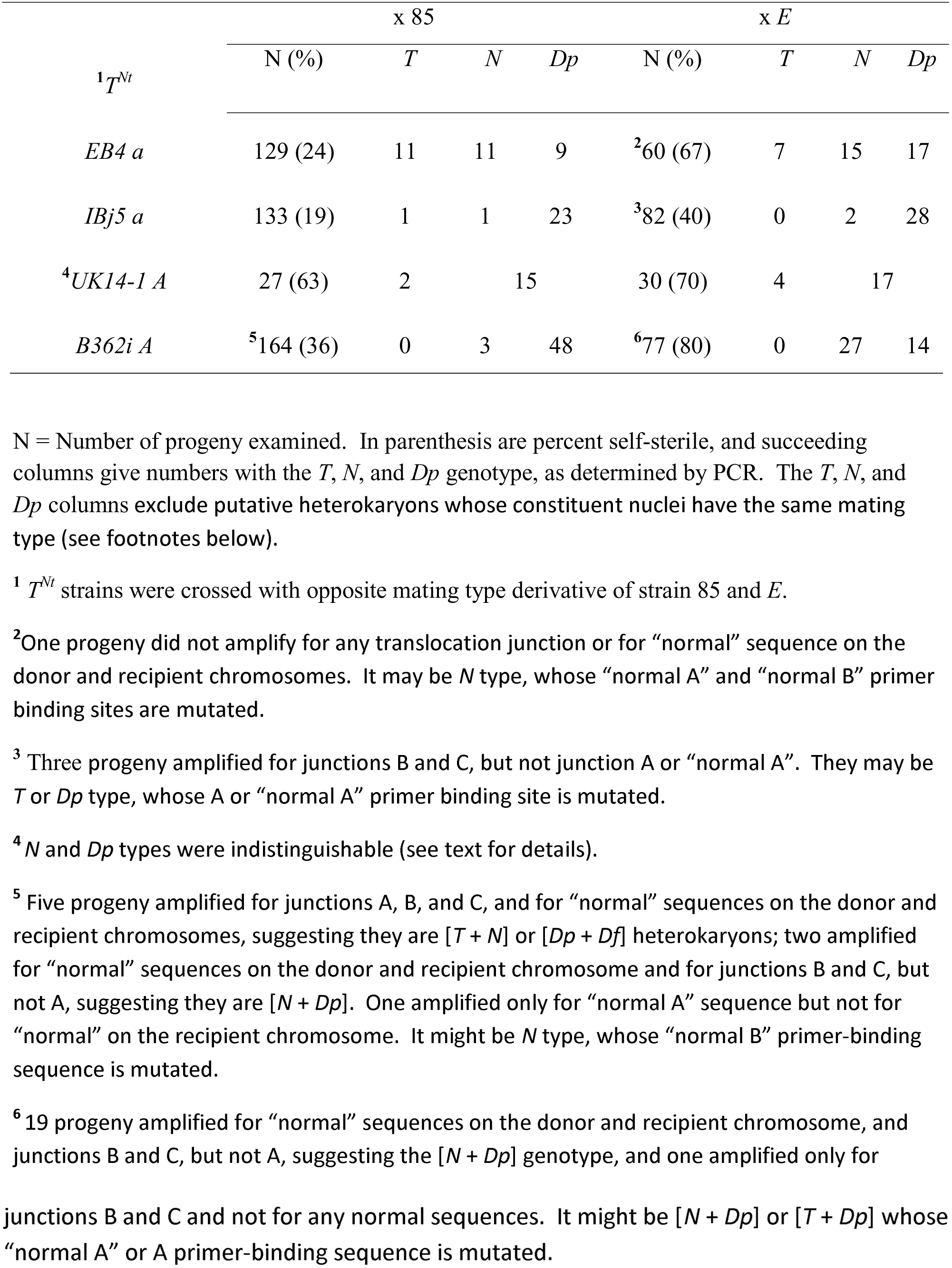
Deviation from the Mendelian ratio in the homokaryotic progeny from *T*^*Nt*^ x *N* crosses.

The crosses of *T(IBj5)*^*Nt*^*a* and *T(B362i)*^*Nt*^*a* with 85 gave *T* and *N* ≪ *Dp*, despite producing 8B:0W asci. This suggested that the black ascospores in the 8B:0W asci might not be as viable as their counterparts from the 4B:4W and 6B:2W asci. To further explore this possibility we harvested ∼500 asci from *T(B362i)*^*Nt*^*A* x 85*a* onto water agar. The majority was four-spored, but we could pick 20 eight-spored asci, including 10 8B:0W, five 6B:2W, four 4B:4W, and one 2B:6W type. In fact, 12 were 7B:1W, 5B:3W, or 3B:5W type, suggesting that in this cross a significant fraction of homokaryotic ascospores remains unpigmented. Table 5 summarizes the number of germinating ascospores from each ascus type. The fraction of black ascospores from 8B:0W asci that produced germlings following the heat-shock to induce germination was lower than that from the 6B:2W and 4B:4W asci.

**Table 5:** Ascospore germination efficiency from 8-spored asci from *T(B362i)*^*Nt*^*A x* 85 *a*

| <sup>1</sup>Asci | N | Black ascospores |  |  |
| --- | --- | --- | --- | --- |
|  |  | P | G | % Germinated |
| 8B:0W | 10 (6) | 56 | 13 | 23 |
| 6B:2W | 5 (4) | 20 | 11 | 55 |
| 4B:4W | 4 (2) | 7 | 3 | 43 |
| 2B:6W | 1 | 1 | 0 | 0 |

<sup>1</sup>Of ~500 asci collected on water agar the majority was four-spored, but 20 were eight-spored. Numbers in parenthesis indicate replacement of 8B:0W asci by 7B:1W; 6B:2W by 5B:3W; and 4B:4W by 3B:5W.
N = number of asci; P = ascospores plated; G = germlings.

Overall, our results provide evidence for deviations from the expected Mendelian ratio among the homokaryotic progeny from crosses of *E* or 85 strains with *T*^*Nt*^ strains which gave the m^-^ phenotype in *T*^*Nt*^ x *E* crosses. The deviations appear to be specific to the homokaryotic progeny because among the heterokaryotic (self-fertile) progeny the [*Dp* + *Df*]/[*T* + *N*] ratios obtained were 3/3 and 7/3 in crosses with *T(IBj5)*^*Nt*^ and *T(B362i)*^*Nt*^ (which showed the m^-^phenotype in crosses with *E*), and 4/4 and 2/3 in crosses with *T(EB4)*^*Nt*^ and *T(UK14-1)*^*Nt*^ (which did not) (Giri *et al*., 2015).

### 3. Introgressing *T*(*VI* > *V*)*UK3-41*

Using genomic DNA of the *T(VR>VIL)UK3-41A* (henceforth, *T(UK3-41) A*) strain as template, and primers complementary to sequences within the translocated segment, we performed inverse PCR (see Materials and methods) and retrieved the adjoining sequence on the recipient chromosome to define the B and C breakpoint junctions (respective Genbank accession numbers KU599833 and KU554720). The inverse PCR for the B junction used genomic DNA digested with *Hae* III, and the flanking sequence had a *Hae* III site located merely 2 bp proximal to the breakpoint junction, which was inadequate to design a primer to PCR amplify across the breakpoint junction. On normal sequence chromosome 6L, a 15882 bp AT-rich (70.5%) sequence containing only a few four-base-pair cutting restriction sites separates the C breakpoint locus from its closest proximal ORF (ncu 07116). These restriction enzymes did not have any convenient sites within the UK3-41 translocated segment, making it difficult to design additional inverse PCR strategies to retrieve additional sequences proximal to the translocated segment. Therefore the B breakpoint remains undetermined. Supplementary Figure 2 updates the breakpoints of 12 *Dp*-generating translocations on the *N. crassa* genome sequence.

Knowing the A and C breakpoint junctions of *T(UK3-41)* allowed us to distinguish the *T*, *Dp* and *N* progeny from *T(UK3-41)* x *N* crosses. Therefore, we attempted to introgress this translocation into *N. tetrasperma* (Figure 2). The initial crosses of *T(UK3-41)* strains with *C4T4 a* and 85 *A* showed phenotype 1 (*T* = *N* = *Dp*), but subsequent to the first productive *T a* x 85 *A* crosses the *T* progeny when crossed with 85 *A* or 85 *a* gave phenotype 3 (*T* ≪ *N* and *Dp*, see above). One introgression line (left in Figure 2), produced no *T* progeny in the f_4_, and another two (middle and right in Figure 2) gave six f_4_ and two f_3_ *T* progeny that were unproductive in subsequent crosses with strain 85 derivatives of opposite mating type. Thus, deviation from the expected Mendelian ratio was one factor that contributed to our inability to introgress this translocation into *N. tetrasperma*.

**Figure 2.**
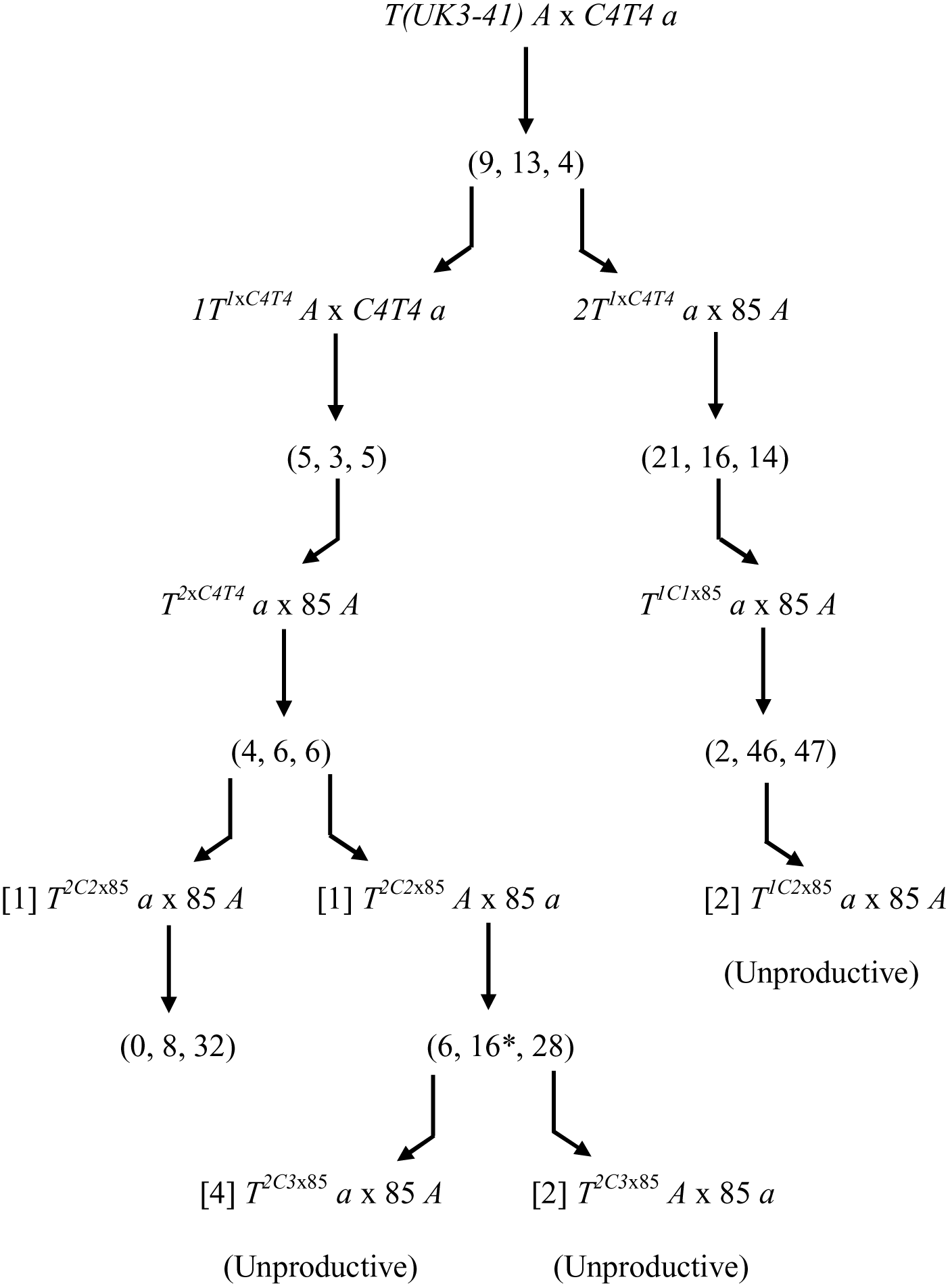
Attempt to introgress *T(UK3-41)* into *N. tetrasperma*. *N. crassa* strain *T(UK3-41) A* was crossed with the *C4T4 a* hybrid strain. PCR with breakpoint junction-specific primers was done to identify the progeny types, and the number of translocation, normal sequence, and duplication types is indicated in the sequence (*T*, *N*, *Dp*). Bent arrows represent the *T* progeny used for the next round of crosses (e.g., *1T*^*1xC4T4*^*A*, *2T*^*1xC4T4*^ *a*). *1T*^*1xC4T4*^*A* x *C4T4 a* yielded the strain *T*^*2xC4T4*^ *a*, which was productive in crosses with *N. tetrasperma* strain 85 *A*. Of the 4 *T*^*2C2*x85^ progeny two were unproductive in crosses, one (*T*^*2C2*x85^ *a*) was productive in the cross with 85 A, and its cross produced 0 *T* progeny, and its sibling strain *2T^1xC4T4^ a* was productive in the cross with *85 A*, producing six *T* progeny strains (4 *T*^*2C3*x85^ *a* and 2 *T*^*2C3*x85^ *A*) that gave unproductive crosses with 85 A or 85 a. One of the 16 *N* type progeny (asterisk) from *T^2C2^*^x85^ *A* x 85 *a* was a self-fertile [*mat A* + *mat a*] heterokaryon. The crosses of two *T*^*1C2*x85^ *a* strains with 85 *A* also were unproductive.

## Discussion

The aim of this study was to try and understand the basis of the striking and unusual max-4^-^ phenotype displayed by some *T*^*Nt*^ x *E* crosses (Giri *et al*., 2015). We still do not know why the *T(IBj5)*^*Nt*^*a* and *T(B362i)*^*Nt*^*A* strains showed the m^-^ phenotype in crosses with *EA*/*Ea* whereas their crosses with 85*A*/85*a* or 2508*A*/2509*a* gave a somewhat weaker phenotype in which a large fraction of ascospores from the 8B:0W asci were inviable. Crosses of the *T*^*Nt*^ strains with the *N. tetrasperma N* strains (85 *A*, 85 *a*, *E A*, *E a*) showed one of three patterns (phenotypes 1, 2, or 3) with respect to transmission ratios in their homokaryotic progeny. In phenotype 1 the *Dp*, *N*, and *T* progeny were produced in comparable numbers. Phenotype 2 yielded mostly (>90%) *Dp* and hardly any *N* or *T* types; and phenotype 3 produced comparable numbers of *Dp* and *N* but the *T* type was absent. Only phenotype 1 is seen in *IT* (and *QT*) x *N* crosses in *N. crassa* (Perkins, 1997; Giri *et al*., 2015), whereas phenotypes 2 and 3 reflect deviations from the expected Mendelian transmission ratio (transmission ratio distortion), and were shown only by crosses with *T*^*Nt*^ strains that induced the m^-^ phenotype in crosses with *E*. The *T* and *N* type progeny are generated by alternate segregation (ALT) in meiosis 1, whereas adjacent-1 segregation (ADJ) produces *Dp* and *Df*. Differential recovery of the products of ALT and ADJ, as in phenotype 2, is a novel type of meiotic drive, one detectable only in crosses of *IT*^*Nt*^ (and *QT*^*Nt*^) with *N*, because ALT and ADJ are not distinguishable in *N* x *N* (and *Dp* x *N*), and all the products from ADJ are inviable in *RT* x *N*. To the best of our knowledge a meiotic drive of comparable nature has been reported in only one other paper: La Chance *et al*. (1964) proposed that ALT was twice as frequent as ADJ in male meiosis heterozygous for a reciprocal translocation in the *Brc* mutant of the screw-worm fly *Cochliomyia hominivorax*. Unfortunately, the Cochliomyia system was not further studied.We do not think ALT is less frequent than ADJ in our system, because it leaves unexplained the production of 0B:8W and 2B:6W asci that are not ordinarily produced in *IT* x *N*.

[*T* + *N*] and [*Dp* + *Df*] asci contain the same genes but distribute them differently in the *mat A* and *mat a* nuclei. Models to explain phenotypes 2 and 3 must translate this difference into differential ascospore viability. One model (model 1) is that the *N. crassa*-derived *T* recipient chromosome and the *N. tetrasperma*-derived *T* donor chromosome homolog carry meiotic drive elements (MDE) that induce inviability in progeny that do not inherit them. MDE are selfish genes that skew the 1:1 Mendelian segregation ratio to their own advantage and their presence, in turn, imposes selection for unlinked suppressors that restore the Mendelian ratio. Presumably, reproductively isolated taxa (e.g., *N. crassa* and *N. tetrasperma*) accumulate distinct drivers and suppressors that get separated in the hybrids, and allow drive to surface in their crosses (Orr *et al*., 2007; Zanders *et al*., 2014). ALT would segregate the MDE into the *T* and *N* progeny thus rendering both inviable, whereas ADJ would segregate both elements into the *Dp* progeny, resulting in their survival. A similar model was proposed to explain the production of more aneuploid or diploid progeny by hybrid *S. kambucha* (*Sk*) / *Schizosaccharomyces pombe* (*Sp*) diploids than even the pure haploid *Sk* and *Sp* parental genotypes (Zanders *et al*., 2014). *Spore killer* elements in Neurospora and Podospora (viz., *Sk-2*, *Sk-3*, *Spok1* and *Spok2*) trigger inviability in ascospores not inheriting them (Turner and Perkins, 1979; Hammond *et al*., 2012; Grognet *et al.*, 2014; Harvey *et al.*, 2014). This model requires that MDE be correctly positioned in *T(IBj5)*^*Nt*^ and *T(B362i)*^*Nt*^. It also does not readily account for phenotype 3. Chromosome 4 is the donor chromosome in *T(B362i)*^*Nt*^, and if an MDE was present on chromosome 4 of the *E* strain, then its segregation in *T(B362i)*^*Nt*^*A* x *E a* could make the *T* progeny that do not inherit it unviable, whereas the *N* and *Dp* progeny, that do, would survive. However, chromosome 4 is the recipient chromosome in *T(IBj5)*^*Nt*^, therefore the model predicts inviability of the *Dp* progeny from *T(IBj5)*^*Nt*^*a* x *E A*, which clearly is not the case.

Model 2 invokes Bateson-Dobzhansky-Muller incompatibility (BDMI) wherein alleles at one locus from one species are unable to function with alleles at another locus from a closely related species (for a short comprehensive review see Presgraves, 2010). BDMIs underlie inviabilty or reduced fertility in several interspecies hybrids (Orr *et al*., 2007; Fishman and Saunders, 2008; McDermott and Noor, 2010; Phadnis, 2011; Zanders *et al*., 2014; Phadnis *et al*., 2015; Sicard *et al*., 2015). BDMI between *N. crassa* and *N. tetrasperma* genes might lead to dysfunction in *T*^*Nt*^ x *N* crosses creating, say, an insufficiency for an ascospore maturation product. Only four viable ascospores (hetero- or homokaryotic) are made following ADJ, whereas four (heterokaryotic) to eight (homokaryotic) viable ascospores can arise following ALT. The generation of >4 viable ascospores might precipitate a “tragedy of the commons”, wherein no ascospore receives a sufficient amount of the maturation factor. This would specifically disfavor the ALT-derived homokaryotic progeny.

How might one explain phenotype 3 (*T* ≪ *N* and *Dp*)? One possibility is that phenotype 2 (*T* and *N* ≪ *Dp*) of the *T(B362i)*^*Nt*^*A* x *E a* cross is masked by the secondary production of homokaryotic *N* cultures via loss of *T* nuclei from heterokaryotic [*T* + *N*] germlings. The *T* genotype has more *N. crassa*-derived sequences than *N*, especially on the translocation donor and recipient chromosomes. Conceivably, a second BDMI selects against the *T* nuclei during vegetative growth. However, we cannot rule out the possibility that the second BDMI affects the viability of the *T* ascospores in [*T* + *N*] asci to produce an effectively four-spored [*T^lethal^* + *N*] ascus. Thus, BDMIs could underlie both phenotypes 2 and 3, and in both cases generate the m^-^phenotype. We would not expect to see any deviation from the Mendelian ratio in crosses with *T*^*Nt*^ strains (e.g., *T(EB4)^Nt^ a*) from which the *N. crassa*-derived MDE (model 1) or BDMI gene(s) (model 2) is absent.

Production of only [*T + N*], [*Dp* + *Df*], and *Dp* progeny can act as a barrier to gene flow from *N. crassa* to *N. tetrasperma*, because the heterokaryotic strains tend not to outcross (Bistis, 1996), and silencing of *Dp*-borne ascus development genes by meiotic silencing by unpaired DNA (MSUD) can render *Dp* x *N* crosses barren (Shiu *et al*. 2001). MSUD is triggered by improper pairing of *Dp*-borne genes in meiosis, causing them to be transcribed into ‘aberrant RNA’, which is processed into MSUD-associated small interfering RNA (masiRNA) used by a silencing complex to identify and degrade complementary mRNA as it exits the nucleus (Hammond *et al*., 2013).

## Acknowledgements

We thank Angela Dheeraj Sharma for assistance with the introgression crosses, and B. Navitha for general technical assistance. DAG was initially supported by a CSIR-UGC Junior Research Fellowship, and later by a CSIR-UGC Senior Research Fellowship. DPK holds the Haldane Chair of the Centre for DNA Fingerprinting and Diagnostics (CDFD). This work was supported by a grant from the Department of Science and Technology, Government of India, and by CDFD Core Funds to DPK.

**Supplementary Figure 1.** Chromosome positions of markers polymorphic between the *N. tetrasperma 85*/*EA*/*Ea* and FGSC 2508*A*/FGSC 2509*a* strains.

**Supplementary Figure 2.** Breakpoints of *Dp*-generating translocations mapped on the *N. crassa* genome sequence. Black lines represent the sequenced chromosomes. Arrowheads indicate the insertion sites of the donor chromosome segments on the recipient chromosomes. Filled arrowheads indicate that the translocated segment is inserted non-inverted relative to the centromere, while open arrowheads signify segments that are inverted relative to the centromere. Open circles indicate genes disrupted by the translocations and filled circles indicate novel open reading frames. The break on the donor chromosome of the quasiterminal translocation *T(UK14-1)* is capped by sequence from the unassigned supercontig 10.9 (star), raising the possibility that supercontig 10.9 might be in distal LG VIL. The figure updates Figure 2 of Singh *et al*. (2010).

